# Arylamine N-acetyltransferase-1 reveals a subpopulation of ALS patients with altered metabolic capacity

**DOI:** 10.1101/2023.06.29.546993

**Authors:** Chandra Choudhury, Sally Allen, Melinder K. Gill, Fleur Garton, Restuadi Restuardi, Neville J. Butcher, Shyuan T. Ngo, Frederik J. Steyn, Rodney F. Minchin

## Abstract

Amyotrophic lateral sclerosis (ALS) is a heterogeneous disease characterised by metabolic changes at onset and throughout disease progression. Here, we investigate the role of arylamine N-acetyltransferase 1 (NAT1), a cytosolic protein associated with mitochondrial function, in ALS. We demonstrate that expression of the murine homolog (mNat2) increases in skeletal muscle of SOD^G93A^ mice, but not control animals, at onset of symptoms and remains elevated until end stage of the disease. Measurement of mitochondrial respiration in peripheral blood mononuclear cells of patients with ALS identified patient sub-populations with low and high metabolic potential, which was strongly associated with NAT1 activity. Those patients with high NAT1 activity had elevated basal respiration, ATP production, mitochondrial reserve, and aerobic glycolysis. NAT1 predicted increased whole body metabolic index, which may be clinically significant as these patients show increased functional decline and shorter survival. NAT1 may be a novel target in those patients with elevated activity.

## Introduction

Amyotrophic lateral sclerosis (ALS) is a progressive neurodegenerative disease associated with the loss of upper and lower motor neurons causing muscle weakness and paralysis. The only currently approved therapies for ALS (riluzole, edaravone and combination sodium phenylbutyrate and taurursodiol) have minimal effects on patient survival (Cho & Shukla, 2020; Miller *et al*, 2012; Paganoni *et al*, 2020) and most succumb to the disease within 3-5 years following diagnosis (Robberecht & Philips, 2013).

Ultrastructural changes in muscle mitochondria from ALS patients were first reported more than 50 years ago (Afifi *et al*, 1966) and since then, altered mitochondrial morphology has been seen in muscle motor neurons (Atsumi, 1981), patient hepatocytes (Nakano *et al*, 1987), and skin cells (Rodríguez *et al*, 2012). Changes in mitochondrial structure have been associated with changes in mitochondrial function (Zhao *et al*, 2022). Mitochondria not only regulate adenosine triphosphate production, but are also central to calcium storage, molecular biosynthesis, and apoptosis (Araujo *et al*, 2020; Pekkurnaz & Wang, 2022). Mitochondrial dysfunction is present in animal models of ALS as well as in patients (Steyn *et al*, 2020), and may be associated with disease severity and survival (Dupuis *et al*, 2011). Aberrant mitochondrial function may contribute to the deterioration in synapses between motor neurons and muscle cells, which can be often seen before any clinical signs of disease (Shefner *et al*, 2023). This has prompted ongoing research to identify pathways that modulate mitochondrial metabolism in ALS (Pekkurnaz & Wang, 2022).

Arylamine N-acetyltransferase 1 (NAT1) is a cytosolic protein associated with mitochondrial function in metabolic diseases such as cancer (Hernández-González *et al*, 2022). Inhibition of NAT1 with small molecule inhibitors or siRNA decreased cell growth (Stepp *et al*, 2018; Tiang *et al*, 2010) and promoted epithelial to mesenchymal transition (Tiang *et al*, 2011). Subsequent studies showed that NAT1 is important in cellular bioenergetics, mitochondrial function, and ATP synthesis (Carlisle *et al*, 2018; Wang *et al*, 2018; Wang *et al*, 2019). NAT1 uses acetyl-coenzyme A as a co-substrate and, in the presence of regulatory molecules such as folate, it can hydrolyse acetyl-coenzyme A to coenzyme A (Laurieri *et al*, 2014). Deletion of NAT1 increases intracellular acetyl-coenzyme A levels (Stepp *et al*, 2019), which are central to mitochondrial respiration. Moreover, ATP, primarily generated in the mitochondria, is an allosteric regulator of NAT1 activity (Minchin *et al*, 2018). NAT1 deficiency inhibits mitochondrial cytochrome C release in response to cytotoxins, resulting in a switch from apoptotic cell death to necroptosis (McAleese *et al*, 2022). While these studies suggest NAT1 is important in mitochondrial function, its role in diseases other than cancer remains largely unexplored.

Here, we hypothesise that altered NAT1 activity in ALS could explain mitochondrial responses to metabolic pressures. To investigate the association between NAT1 activity and mitochondrial function, we examined NAT1 expression and activity in the SOD1^G93A^ mouse model of ALS as well as peripheral blood mononuclear cells (PBMC) isolated from patients with ALS. We report that NAT1 expression identifies a sub-population of patients with elevated mitochondrial function that is not seen in healthy individuals and suggests NAT1 may be a novel target in these patients.

## Results

### NAT1 is increased early and persistently throughout the course of disease in the SOD1^G93A^ mouse model of ALS

SOD1^G93A^ mice recapitulate many of the symptoms and phenotypes seen in patients with ALS (Philips & Rothstein, 2015). Disease progression in SOD1^G93A^ mice is defined by 4 stages: pre-onset, onset, mid-stage, and end-stage of disease (Fig 1A). Disease onset includes the loss of spinal motor neurons and denervation of muscle, generally observed at 50-60 days of age. This worsens with disease progression (Matsumoto *et al*, 2006), and is associated with weight loss. Changes in mitochondrial function have been observed in SOD1^G93A^ mouse neurons and skeletal muscle prior to the onset of disease (Pharaoh *et al*, 2019; Scaricamazza *et al*, 2020).

**Figure. 1.**
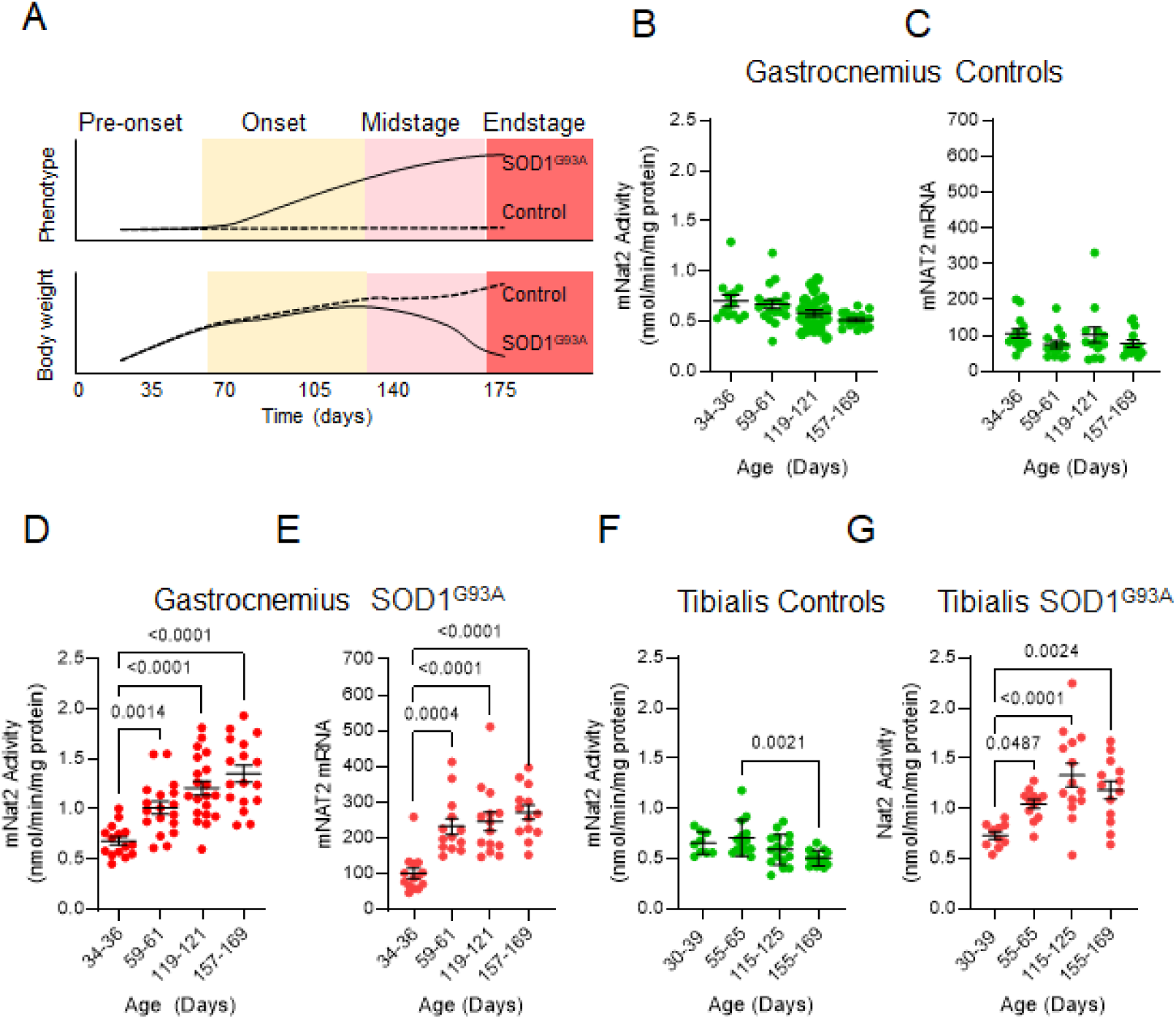
Expression of mNat2 in SOD1^G93A^ mouse tissues. A Disease progression in SOD1^G93A^ mice with approximate timeline throughout life. Pre-onset mice present with a lack of phenotype and demonstrate normal body weight gain when compared to controls. Disease onset is associated with the emergence of a phenotype defined by fine motor tremors in the hindlimbs. Changes in locomotive velocity, rearing episodes and paw grip endurance, and weight loss are common in mid-stage disease. At end-stage, mice have limited movement and are unable to right themselves within 15 sec. B mNat2 activity and in gastrocnemius muscle from control animals culled at different ages corresponding to disease stages observed in SOD1^G93A^ mice (mean ± s.e.m, n = 13-24 animals). C mNat2 mRNA levels in gastrocnemius muscle from control animals (mean ± s.e.m, n = 13-14 animals). D mNat2 activity in gastrocnemius muscle from SOD1^G93A^ mice at ages corresponding to the different stages of disease (mean ± s.e.m, n = 15-20 animals, ANOVA with Bonferroni correction). E mNat2 mRNA in gastrocnemius muscle from SOD1^G93A^ mice at the different ages corresponding to the stages of disease (mean ± s.e.m, n = 13-14 animals, ANOVA with Bonferroni correction). F mNat2 activity in tibialis anterior muscle from control animals culled at different ages corresponding to the stages of disease in SOD1^G93A^ mice (mean ± s.e.m, n = 9-16 animals). G mNat2 activity in SOD1^G93A^ mice at different ages corresponding to the stages of disease (mean ± s.e.m, n = 10-14 animals, ANOVA with Bonferroni correction).

Since human NAT1 is associated with mitochondrial function, we assessed the activity of murine Nat2 (mNat2; the homolog of human NAT1) in the gastrocnemius muscle of SOD1^G93A^ mice as disease progressed. This muscle comprises a mixture of fast and slow twitch fibres. Neither mNat2 activity (Fig 1B) nor mNat2 mRNA (Fig 1C) changed with age in control mice. In SOD1^G93A^ mice, activity was similar to controls prior to disease onset (Fig 1D). However, at an age corresponding to disease onset, mNat2 activity (Fig 1 D) and mRNA expression (Fig 1E) increased. This increase was maintained through to disease end-stage. We also examined mNat2 activity in the tibialis anterior muscle from control (Fig 1F) and SOD1^G93A^ mice (Fig 1G), a muscle that primarily comprises fast twitch fibres. There was no change in activity with age in the control mice (Fig 1F). As with the gastrocnemius muscle, mNat2 increased at an age corresponding to disease onset in SOD1^G93A^ mice. This increase in expression was sustained throughout disease course (Fig 1G). Collectively, the results show an early and persistent increase in NAT1 activity in muscle from ALS mice.

### Mitochondrial respiration and glycolysis are altered in ALS PBMC

We next asked whether there are differences in cellular metabolic processes between patients with ALS and non-neurodegenerative disease controls, and whether these changes are related to NAT1 expression. To do this, we examined NAT1 activity and metabolic function in patient and control derived PBMC. Prior studies have shown mitochondrial changes in ALS PBMC, including changes in gene expression (Luotti *et al*, 2020), mitophagy (Bordoni *et al*, 2022) and mitochondrial respiration (Araujo *et al*., 2020).

We compared mitochondrial oxidative phosphorylation and aerobic glycolysis in PBMC collected from 33 patients with ALS (Cases) and 24 healthy non-neurodegenerative disease controls (Controls) (Table 1). Controls and Cases were matched by age, body mass index (BMI) and fat free mass. While there was a slight difference in sex distribution, this was not significant (Chi-squared with Yates correction, p = 0.37). Basal oxygen consumption, ATP production, reserve respiratory capacity (RRC), and maximal respiration in whole cells were quantified by monitoring oxygen consumption following the sequential addition of oligomycin, FCCP and rotenone/antimycin A, respectively (Fig 2A). Mitochondrial protein leak and non-mitochondrial respiration were also determined. There were no significant differences in the mean basal respiration (Fig 2B) or ATP production (Fig 2C) in PBMC from Controls versus Cases (Mann-Whitney test, p > 0.05) although the skewness (1.4-1.5) and excess kurtosis (2.3-3.0) indicated that the patient data were asymmetric having a larger number of outliers with higher values compared to Controls. This was confirmed by testing for normality using the D’Agostino and Pearson Test and the Shapiro-Wilk Test (Supplementary Table S1). Similar differences in data distribution were seen for RRC (Fig 2D) and maximal respiration (Fig 2E), although there were no differences in the means. Neither protein leak (Fig 2F) nor non-mitochondrial respiration (Fig 2g) differed between the two groups.

**Fig 2.**
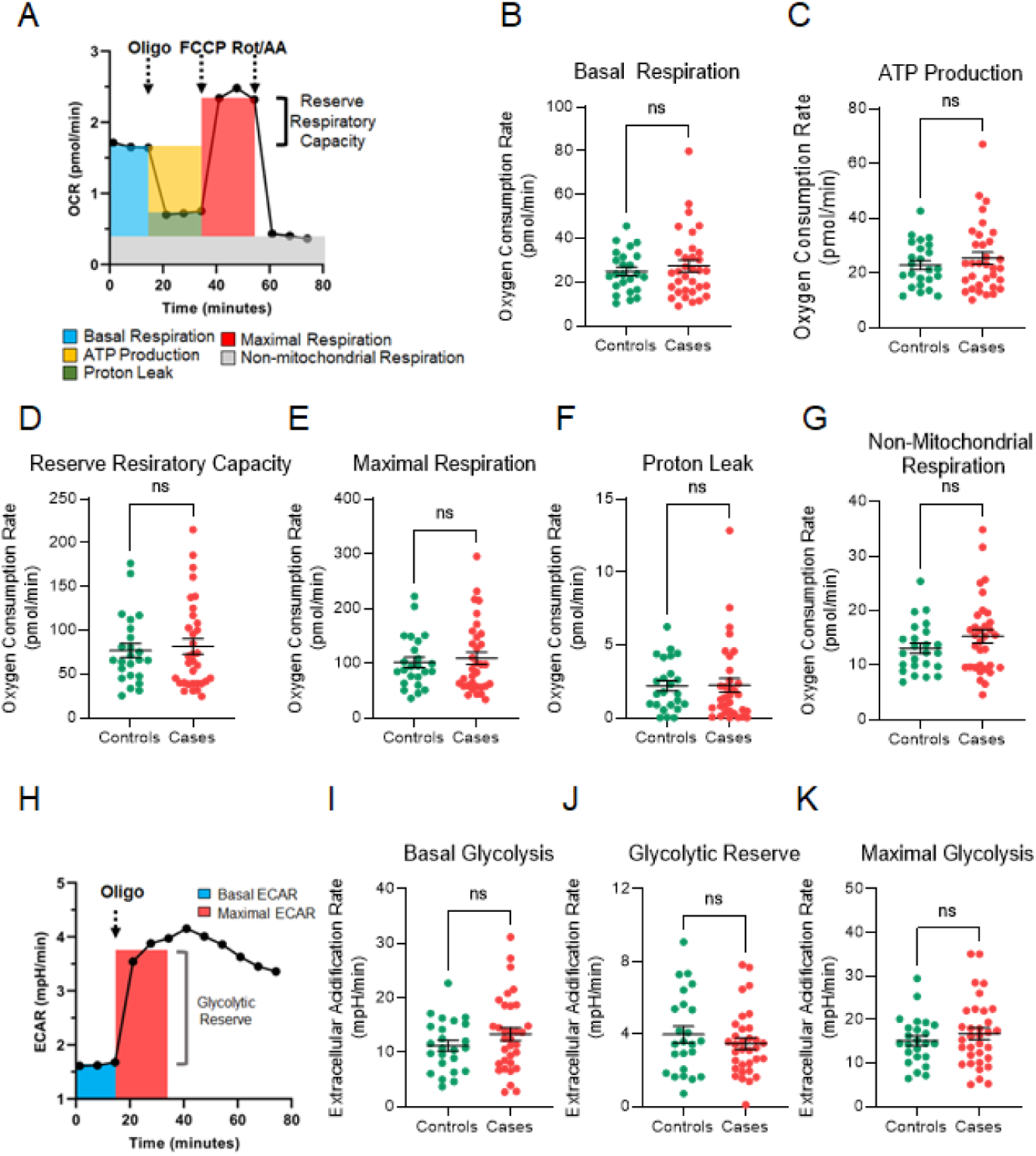
Mitochondrial respiration and glycolysis in PBMC isolated from Controls and patients with ALS. A Schematic representation for quantification of mitochondrial oxygen consumption rate (OCR) using specific inhibitors of respiration. Oligomycin (Oligo) inhibits ATP synthase, FCCP disrupts the mitochondrial membrane potential, rotenone (Rot) and antimycin A (AA) inhibits total mitochondrial respiration. B Basal respiration in non-neurodegenerative disease controls (Controls, n = 24) and patients with ALS (Cases, n = 33). C ATP production in Controls and Cases D Reserve respiratory capacity in Controls and Cases E Maximal respiration in Controls and Cases F Proton leak in Controls and Cases G Non-mitochondrial respiration in Controls and Cases. H Schematic representation for the quantification of aerobic glycolysis using extracellular acidification rate (ECAR) following the inhibition of mitochondrial function with oligomycin. I Basal glycolysis in Controls and Cases J Glycolytic reserve in Controls and Cases K Maximal glycolysis in Controls and Cases. All data are the mean of quadruplicate measurements for each individual. Comparisons were performed using a Mann-Whitney test. ns = not significant. All data are shown as mean ± s.e.m.

**Table 1.** Demographics of Participants at Time of Assessment.

|  | Controls | Cases (Activity cohort) |
| --- | --- | --- |
| N | 24 | 33 |
| Age in years (SD) | 56.4 ± 9.5 | 60.4 ± 9.7 |
| Sex | M = 14 (58%) F = 10 (42%) | M = 23 (70%) F = 10 (30%) |
| BMI (SD) | 30.6 ± 9.0 | 26.5 ± 5.2 |
| % Free Fat Mass (SD.) | 62.5 ± 2.1 | 64.4 ± 1.8 |
| Riluzole Treatment |  | 22 (60%) |
| Age of Onset years (SD) |  | 56.6 ± 9.9 |

SD = standard deviation, BMI = body mass index

Aerobic glycolysis generates an increase in extracellular acidification during the conversion of glucose to pyruvate and is maximal when mitochondrial respiration is inhibited. To quantify glycolysis, the extracellular acidification rate was measured in PBMCs before and after the addition of oligomycin, an inhibitor of mitochondrial ATP synthase (Fig 2H). Neither basal glycolysis (Fig 2I), glycolytic reserve (Fig 2J) or maximal glycolysis (Fig 2K) differed between cases and controls. Moreover, the skewness was similar between the different groups (Supplementary Table S1).

Collectively, these results show little difference in the mean values for mitochondrial respiration or aerobic glycolysis in PBMCs from Controls and patients with ALS. However, the distribution of some parameters associated with mitochondrial respiration was normal in Controls but non-normal in the Cases. While the reason for this difference is not known, one explanation is the presence of greater heterogeneity in ALS due to the disease. This may be significant as identification of sub-populations in other diseases has had a major impact on treatment outcomes with targeted personalised medicine (Yamamoto *et al*, 2022). Detecting clinically relevant sub-populations in ALS has been challenging. To address the possibility of sub-populations in ALS, we further examined the metabolic profile of Controls and Cases.

### PBMC metabolic phenotypes reveal sub-populations of ALS patients

Basal bioenergetics were analysed by plotting basal extracellular acidification rates against basal oxygen consumption rates (Fig 3A and B). Each plot was divided into quadrants that represented the phenotypes of quiescent, glycolytic, oxidative, and energetic metabolism in the resting state. There was a strong positive correlation with both Controls (Spearman rho (ρ) = 0.85, p < 0.001) and Cases (ρ = 0.91, p < 0.001). However, most of the Controls (92%) showed a quiescent phenotype characterised by low rates of oxidative phosphorylation and glycolysis (Fig 3a; lower left quadrant). By comparison, 26% of patient derived PBMCs showed a glycolytic or energetic phenotype (Fig 3B; upper right quadrant), indicating PBMCs from these individuals have higher oxidative phosphorylation and aerobic glycolysis when compared to the other Cases and the Controls. Maximal acidification rates and oxygen consumption rates measure the capacity for mitochondria to function under metabolic stress. When these were plotted, approximately 10% of controls (Fig 3C) and 30% of patients (Fig 3D) had an energetic phenotype.

**Fig. 3.**
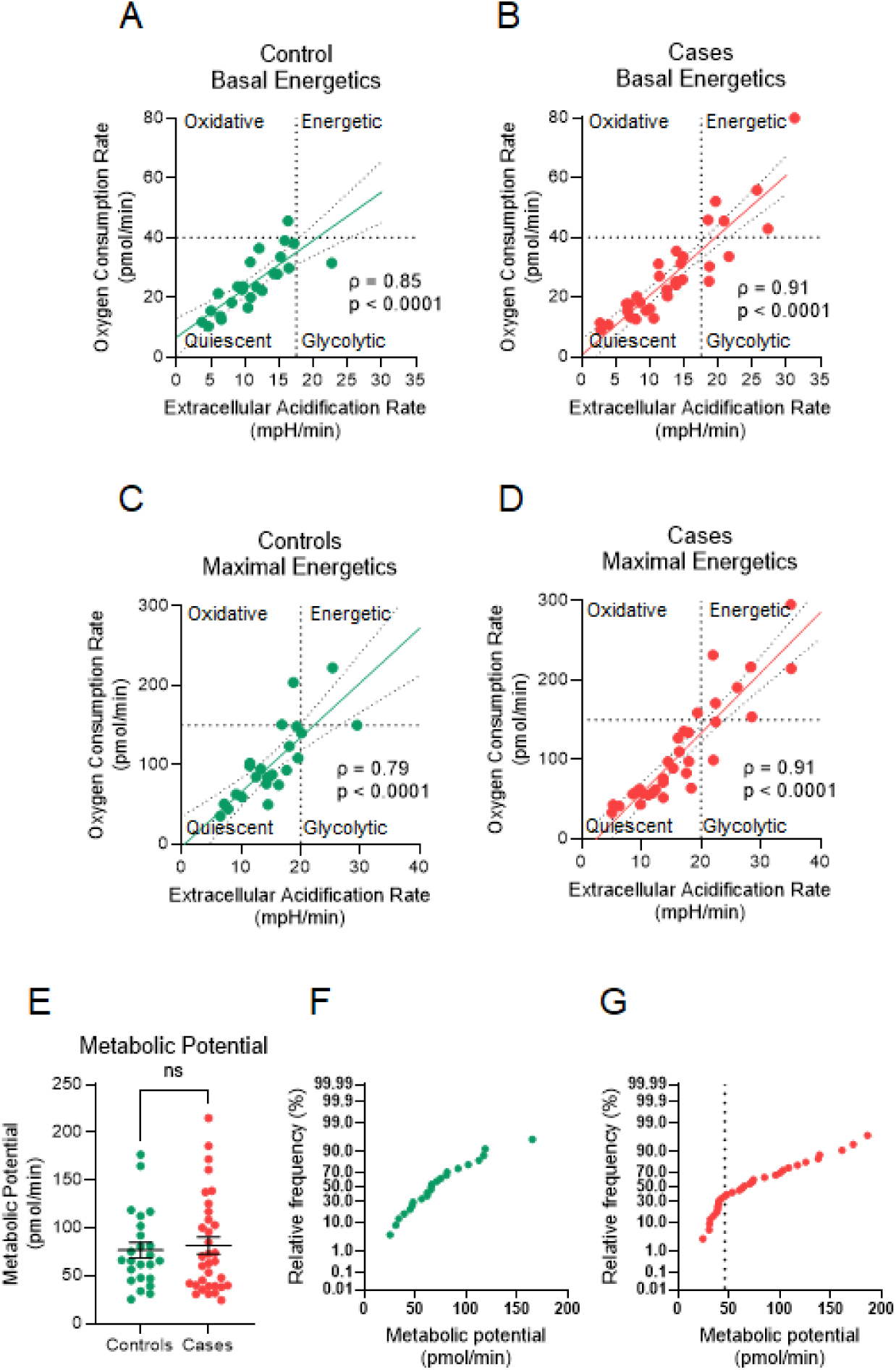
PBMC energy phenotype in sub-populations of ALS patients. A Plot of basal extracellular acidification rate versus basal oxygen consumption rate (basal energetics) in control individuals (n = 24). The plot was divided into quadrants that represented the different bioenergetic states of the cell. Solid lines represent simple linear regression with 95% confidence limits (dotted lines). B Basal energetics for patients with ALS (Cases, n = 33). C Maximal energetics for Controls. D Maximal energetics for Cases. E Metabolic potential was calculated as the sum of the mitochondrial reserve respiratory capacity and the glycolytic reserve and compared using Mann-Whitney U-test. ns = not significant. Data are mean ± s.e.m. F Frequency distribution (Probit plot) of metabolic potential in the control population. G Frequency distribution (Probit plot) of metabolic potential in cases shows two populations with a cut-off of ∼48 pmol/min (dotted line).

Metabolic potential is the difference between basal and maximal respiration and is a measure of the spare capacity of a cell to increase metabolism when required. Cells with a low metabolic potential respond less to cues for increased energy demand compared to those with a high metabolic potential (de Jong *et al*, 2022). For both controls and patients, there was no significant difference between the mean values for metabolic potential (Fig 3E). However, the distributions of the data showed interesting distinctions. For the controls, a cumulative frequency plot of metabolic potential was curvilinear suggesting the presence of a single population (Fig 3F). By contrast, the same plot for the ALS patients (Fig 3G) showed a distinct sub-population (metabolic potential < 48 pmol/min) where the relative frequencies deviated from a line of best fit. This sub-population, which represented 37% of all cases, had the lowest capacity to respond to metabolic stress.

### Low metabolic potential is associated with high body mass

Cases and Controls were divided into those with low metabolic potential and those with high metabolic potential based on the cut-off of 48 pmol/min (see Fig 3G). RRC was significantly elevated in participants with high metabolic potential in both controls and in patients (Fig 4A), which was expected since this parameter is the major determinant of metabolic potential. Glycolytic reserve was not different between the two groups in the controls but was greater in the cases with high metabolic potential (Fig 4B). Other measures of mitochondrial respiration were also different between the two groups. Basal respiration (Fig 4C) and basal glycolysis Fig 4D) were greater in those with high metabolic potential, for both controls and cases. These results indicate lower overall mitochondrial bioenergetics in individuals with low spare metabolic capacity, both in the resting (basal) and stressed (RRC) state.

**Fig. 4.**
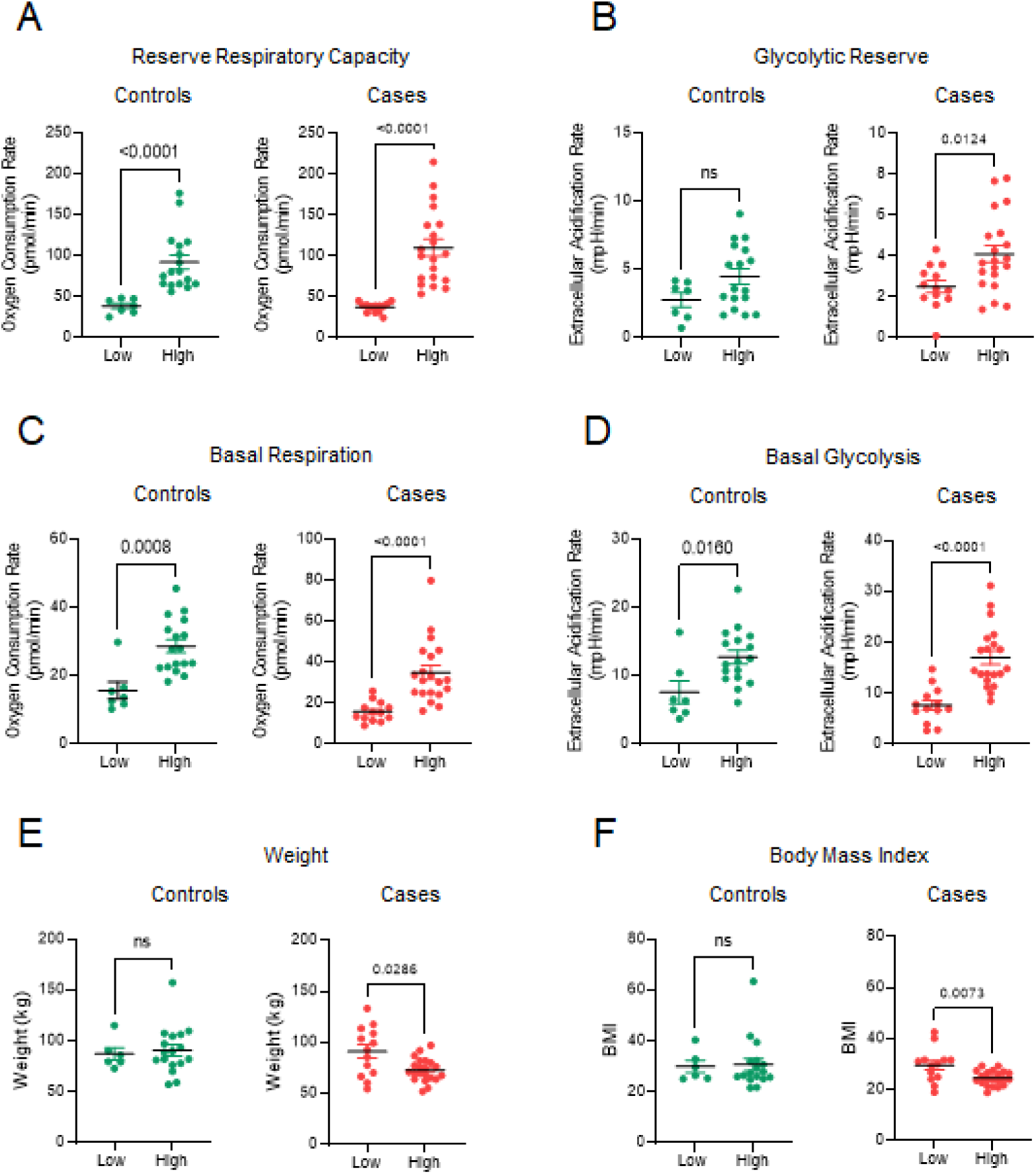
Low metabolic potential is associated with high body mass index. Controls and Cases were divided into low and high metabolic potential groups using a cut-off of 48 pmol/min (as defined in Fig 3g). A Reserve respiratory capacity in Controls and Cases. B Glycolytic reserve in Controls and Cases. C Basal respiration in Controls and Cases. D Basal glycolysis in Controls and Cases. E Body weight in Controls and Cases. F Body mass index in controls and cases. All data are shown as mean ± s.e.m. and compared using a Mann-Whitney test.

Increased body mass is related to low mitochondrial respiration, especially in adipose tissue (Yin *et al*, 2014) and skeletal muscle (Bakkman *et al*, 2010). Consequently, we investigated whether body mass varied with metabolic potential. In control subjects, there was no difference whereas patients with high spare metabolic capacity had a significantly lower body mass (Fig 4E). A similar result was seen for BMI in both cohorts (Fig 4F). The difference in overall body mass was due to a higher fat mass in the low metabolic potential individuals (low = 36.9 ± 4.5 kg versus high = 24.8 ± 1.9 kg, p < 0.01, Mann-Whitney test). Interestingly, fat mass in the low metabolic potential patients was similar to that seen in the controls (33.9 ± 2.8 kg) indicating that patients, but not controls, with elevated spare metabolic potential were leaner.

### NAT1 activity in PBMC from ALS patients

To determine whether up-regulation of NAT1 is seen in patients with ALS, we measured NAT1 activity in PBMCs. There was a significant increase in NAT1 in patients similar to that seen in mouse muscle tissues (Fig 5A). To further validate this finding, we examined published NAT1 expression data that quantified mRNA isolated from blood cells. The dataset (GSE112680) comprised 137 controls and 164 cases (Swindell *et al*, 2019). Similar to our own data, NAT1 expression was higher in the Cases when compared to Controls (Fig 5B).

**Fig. 5.**
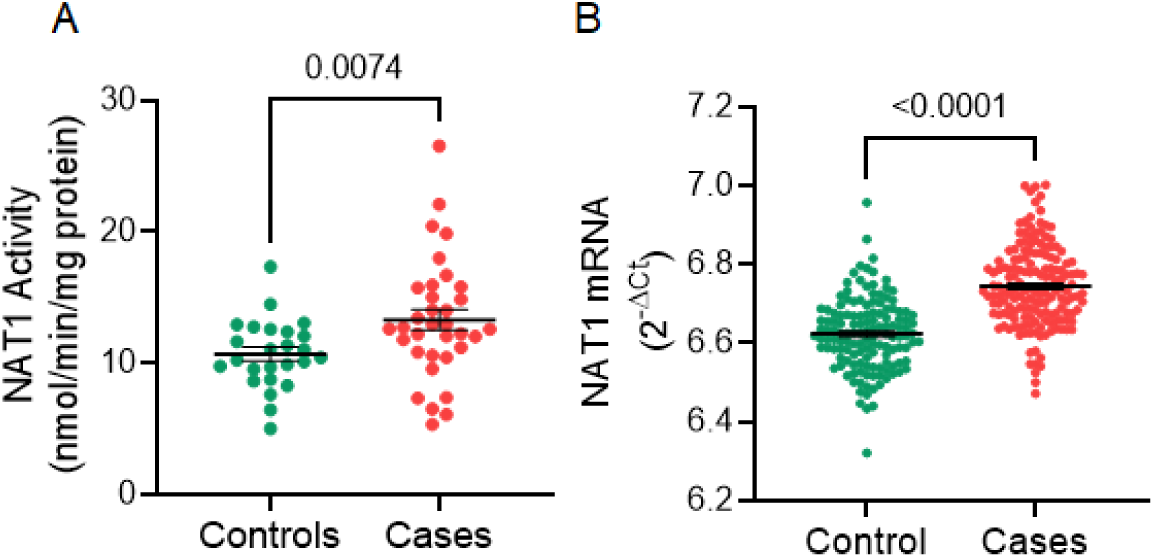
NAT1 expression is elevated in ALS patients. A NAT1 activity in controls (n = 24) and cases (n = 33). B Published data from Swindell et al (Swindell *et al*., 2019) that included controls (n = 137) and cases (n = 164). Groups were compared using a Mann-Whitney U test, with p value shown. Data are mean ± s.e.m.

### Genetic variation does not explain elevated NAT1 activity in ALS patients

The *NAT1* gene is genetically variant, with polymorphisms in both the open reading frame and the 3’UTR that affect protein stability and enzyme activity (Hein *et al*, 2000). To investigate whether *NAT1* polymorphisms explain the differences in NAT1 activities, DNA isolated from PBMCs was sequenced across the coding region and the 3’UTR of the *NAT1* gene. Single nucleotide polymorphisms (SNPs) were then used to deduce *NAT1* haplotypes according to the Arylamine N-acetyltransferase Nomenclature database (http://nat.mbg.duth.gr/). The currently used reference allele (*NAT1*4*) accounted for 64% and 71% of all alleles in the controls and cases, respectively (Table 2). However, there were no differences in allele distribution between the two groups (Chi Squared, p>0.05). These results suggest that genetic polymorphisms do not account for the differences in NAT1 expression between Controls and Cases.

**Table 2.** NAT1 haplotypes in controls and cases.

| Allele | Effect on Activity | Phenotype | Frequency (%) |  |
| --- | --- | --- | --- | --- |
|  |  |  | Controls (n = 44) | Cases (n = 58) |
| NAT1*4 (ref) | None | Intermediate | 28 (64%) | 41 (71%) |
| NAT1*10 | Increase | Rapid | 10 (36%) | 12 (21%) |
| NAT1*14A & B | Decrease | Slow | 2 (4%) | 2 (3%) |
| NAT1*18A | None | Intermediate | 1 (2%) | 0 |
| NAT1*26B | None | Intermediate | 1 (2%) | 0 |
| NAT1*17 | Decrease | Slow | 2 (4%) | 1 (1%) |
| NAT1*11B | None | Intermediate | 0 | 1 (1%) |
| NAT1*3 | None | Intermediate | 0 | 1 (1%) |

NAT1\*4 was used as the reference gene (ref). Effects of each allele on enzyme activity was obtained from the Arylamine N-acetyltransferase Nomenclature database and references therein (http://nat.mbg.duth.gr). Total number of alleles (n) in the cases and controls is shown in parentheses.

We further interrogated a recently published ALS GWAS database containing SNPs from 22,205 ALS patients and 110,881 controls (van Rheenen *et al*, 2021) for the frequency of 17 different SNPs associated with NAT1 phenotypes, of which 11 were found in the database (Supplementary Table S2). None of the SNPs were associated with ALS (p-value range = 0.32 – 0.93). Finally, to determine whether other SNPs located near NAT1 may differ between Cases and Controls, eQTL analysis was performed to identify further cis-acting SNPs (within 1.5 Mb) that alter NAT1 expression. Of the 45 independent SNPs extracted from the GTEx database, none met the p-value threshold of 0.00075 (0.05 with Bonferroni correction).

From the patient data in the current study as well as results from a large database, there is no apparent difference in the genetic variation of the NAT1 gene between Controls and patients with ALS, suggesting that other factors such as epigenetic differences account for the high NAT1 activity seen in a sub-population of ALS patients.

### NAT1 activity and cell bioenergetics in PBMC from ALS patients

We found that PBMC from some ALS patients have higher basal as well as maximal bioenergetics compared to Controls (Fig 3), suggesting these individuals have an advantage during metabolic stress. To determine whether NAT1 activity was associated with the capacity for PBMC to respond to stress, activity was plotted against metabolic potential. There was no association in Controls (Fig 6A) but a significant positive association in PBMC isolated from patients with ALS (Fig 6B). Moreover, in approximately 30% of cases, NAT1 activity exceeded that seen in the Controls.

**Fig. 6.**
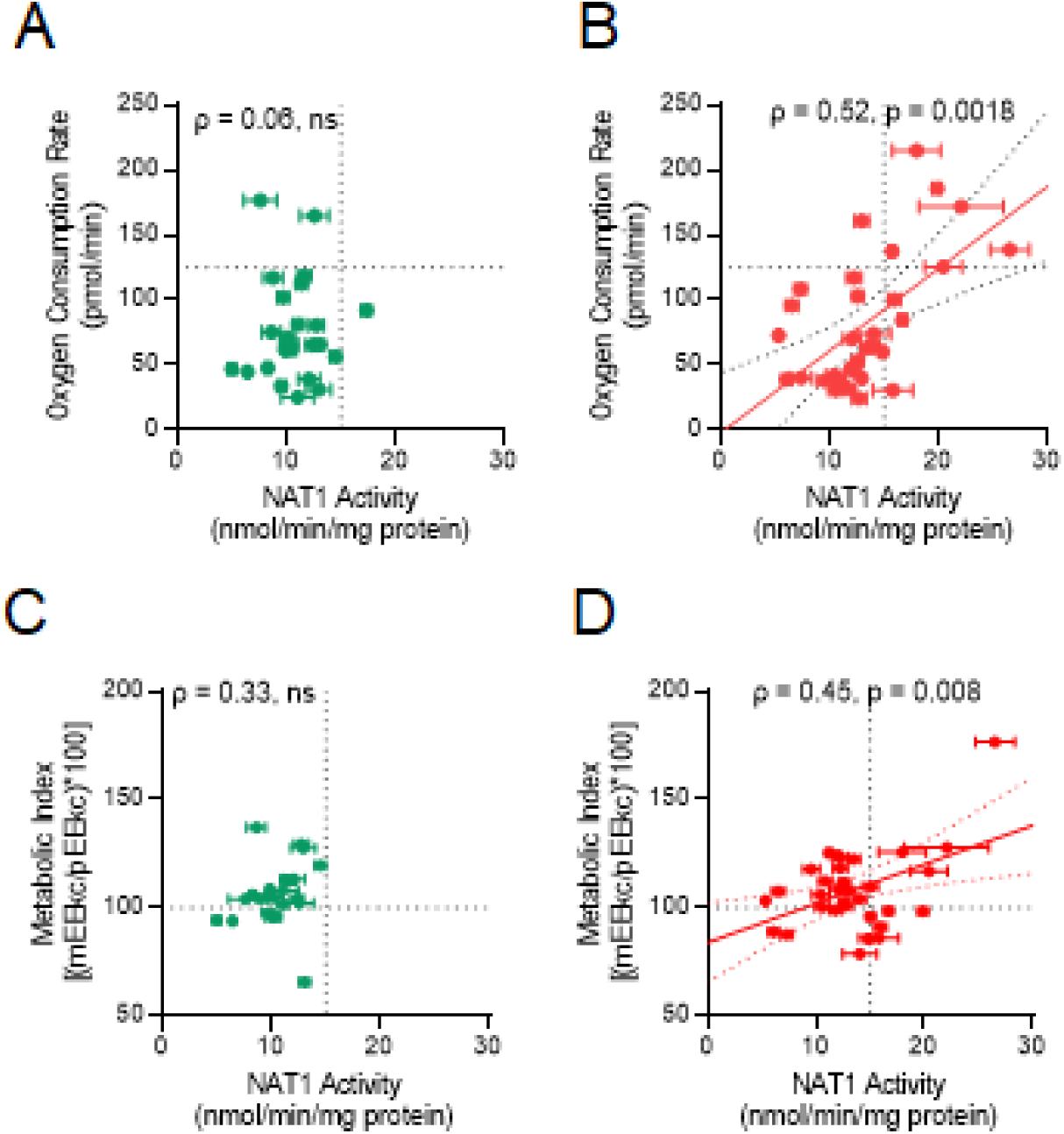
High NAT1 activity in ALS patients is associated with increased metabolic potential. A NAT1 activity (mean ± s.e.m, n = 3 independent observations for each subject) versus metabolic potential in Controls. There was no association between the two variables (Spearman’s ρ = 0.06). B NAT1 activity versus metabolic potential in ALS patients (Cases). Spearman’s ρ = 0.52 (p = 0.018). Linear regression of the data is shown by the solid line with the 95% confidence interval shown by the dotted lines. C NAT1 activity versus metabolic index in controls. There was no association (Spearman’s ρ = 0.33). D NAT1 activity versus metabolic index in Cases. Spearman’s ρ = 0.45, p =0.008. Linear regression of the data is shown by the solid line with the 95% confidence interval shown by the dotted lines.

We assessed whole body energy expenditure for each individual on the same day as blood collection for isolation and assessment of NAT1activity in PBMC. Metabolic index (MI) is a ratio of measured energy expenditure relative to expected energy expenditure and has been used previously to identify hypermetabolic individuals (Steyn *et al*, 2018; Steyn *et al*., 2020). We found no association between metabolic index and NAT1 activity in Controls (Fig 6C) whereas there was a strong positive correlation in patients with ALS (ρ = 0.45, p = 0.008; Fig 6d. This observation may be important clinically because some patients with an elevated MI show increased functional decline and shorter survival (Steyn *et al*., 2018).

### PBMC NAT1 activity is associated with mitochondrial function in patients with ALS, but not in Controls

To determine whether NAT1 expression was associated with cellular respiration, NAT1 activity was plotted against basal respiration (Fig 7A), ATP production (Fig 7B), RRC (Fig 7C) and maximal respiration (Fig 7D). In Controls, there was no association suggesting NAT1 expression does not contribute to the regulation of mitochondrial respiration in NORMAL PBMCs. By contrast, mitochondrial respiration and NAT1 activities were positively correlated in patients with ALS. This was observed for basal respiration, ATP production, RRC and maximal respiration. A similar difference between Controls and Cases was also seen for non-mitochondrial respiration (glycolysis), where a positive correlation was observed in ALS patients for basal glycolysis (Fig 7E) and maximal glycolysis (Fig 7F). Again, no correlations were observed in the controls. The positive associations in the Cases were driven mostly by patients with NAT1 activities greater than 15 nmol/min/mg protein, suggesting these individuals have disease-associated mitochondrial hyperactivity, with higher basal respiration, increased ATP production and greater reserve.

**Fig. 7.**
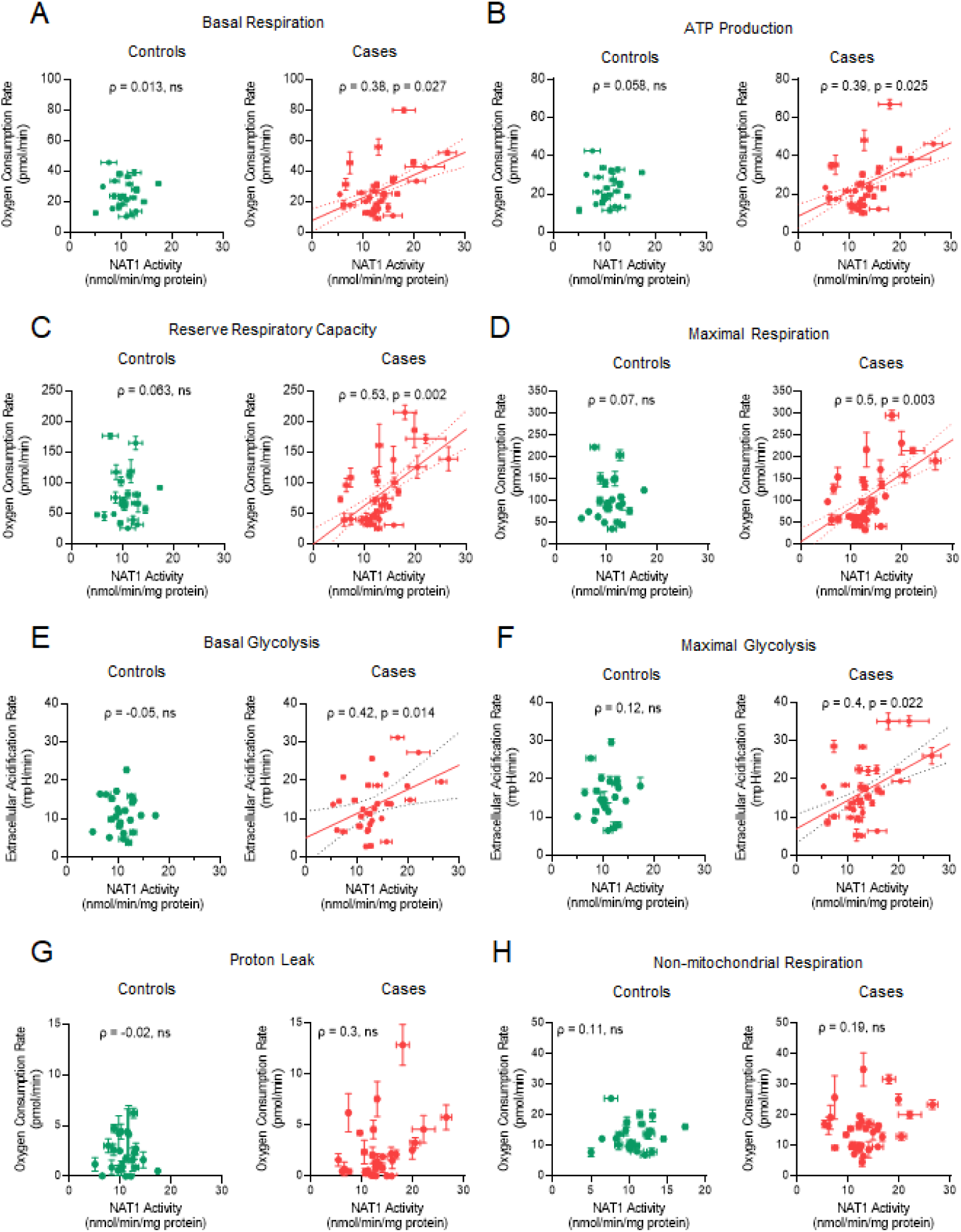
Association between NAT1 activity and cellular respiration. A Basal respiration, B ATP production, C Reserve respiratory capacity, D Maximal respiration, E Basal glycolysis F Maximal glycolysis in Controls and Cases. Associations were computed using Spearman’s rank correlation with rho (ρ) shown for each graph. Where ρ < 0.05, simple linear regression was also computed and is shown with 95% confidence limits (dotted lines). ns = not significant (p > 0.05). In each graph, NAT1 activity = mean ± s.e.m, n = 3.

## Discussion

SOD1^G93A^ mice are arguably the most studied mouse model of human ALS but, unlike the human disease, they are genetically homogeneous with well-defined predictable phenotypes. Disease onset occurs at approximately 50 days of age where loss of spinal motor neurons and the denervation of muscle is observed. This is also associated with the emergence of metabolic phenotypes, including weight loss. Recent work showed fatal ALS-like disease in mice where expression of SOD^G93A^ was restricted to skeletal muscle (Martin & Wong, 2020), suggesting that metabolic changes in muscle may not be secondary to changes in the CNS, at least in the SOD^G93A^ mouse model. We found that, at the time of disease onset, mNat2 increased in a disease-dependent manner in SOD^G93A^ mice. This was observed in gastrocnemius and tibialis muscle and involved both mNat2 mRNA and enzyme activity. In human PBMCs, a similar increase in the activity of the human homolog of mNat2 (NAT1) was observed, albeit only in a sub-population of patients.

Surrogate markers of ALS in PBMC have been previously reported, including the presence of aggregated SOD1 (Pansarasa *et al*, 2022). Moreover, in a study involving 93 patients and 104 controls, Luotti and colleagues showed that decreased expression of peptidyl-prolyl cis-trans isomerase A in PBMC is a disease modifier and predicted rapid disease progression (Luotti *et al*.). PBMCs from patients with ALS have decreased mitochondrial calcium uptake and increased autophagy-related gene expression (Araujo *et al*., 2020). PBMCs also show changes in mitochondrial respiration in patients with metabolic dysfunction such as end-stage kidney disease (Altintas *et al*, 2021), inborn errors of fatty acid oxidation (Stenlid *et al*, 2022), hypertension (Sommer *et al*, 2022) and heart failure (Li *et al*, 2015). Thus, PBMCs have emerged as a reliable proxy for assessing metabolic perturbations in diseases. In the present study, PBMCs failed to show any differences in mean mitochondrial respiration between Cases and Controls. However, previous studies have demonstrated other changes in mitochondrial function such as calcium uptake and redox homeostasis (Araujo *et al*., 2020). We did observe greater metabolic heterogeneity in PBMCs within patients with ALS, but not controls, with a larger proportion of patient PBMCs having a glycolytic/energetic phenotype. Although the patient number was small (n = 7) Echaniz-Laguna *et al* found that maximal mitochondrial respiration in muscle biopsies increases with disease progression while there were no changes in basal respiration (Echaniz-Laguna *et al*, 2006). In a larger study (n = 50), loss of cytochrome C oxidase was seen in skeletal muscle biopsies from ALS patients, although a high degree of inter-patient heterogeneity was evident (Crugnola *et al*, 2010). Collectively, these results suggest that mitochondrial dysfunction in ALS is common, but the extent of perturbation varies between patients.

We identified a group of ALS patients with low metabolic potential in PBMCs. These individuals had a significantly higher BMI and a 50% increase in fat mass. Metabolic potential is primarily determined by RRC, which is a measure of how much additional ATP can be synthesised by mitochondria when energy demands increase. Healthy mitochondria have a high RRC whereas aging or dysfunctional mitochondria may have little or none (Marchetti *et al*, 2020). RCC is regulated by the supply of mitochondrial substrates, in particular glucose-derived pyruvate. Inhibition of the pyruvate dehydrogenase complex, which is essential for pyruvate utilisation by mitochondria, diminishes RRC (Marchetti *et al*., 2020). The patients with low RRC also had the lowest NAT1 activities (Fig 6B). It is noteworthy that NAT1 deficiency in cancer cells decreases pyruvate dehydrogenase activity, which can be overcome by inhibiting pyruvate dehydrogenase kinase (Wang *et al*., 2019). Inhibitors of pyruvate dehydrogenase kinase have been studied in ALS. For example, dichloroacetate reduces disease pathology in SOD1^G93A^ mice (Martínez-Palma *et al*, 2019) which is consistent with the reported up-regulation of pyruvate dehydrogenase kinase 4 in skeletal muscle from these animals following disease onset (Palamiuc *et al*, 2015). Pyruvate dehydrogenase kinase inhibitors have not progressed to trials in ALS patients, although such treatments might be most beneficial to those patients with low RRC.

In the absence of sufficient pyruvate supply to the TCA cycle, mitochondria switch to fatty acid oxidation to maintain RRC (Marchetti *et al*., 2020). Increased fatty acid dependency in skeletal muscle has been reported for both SOD1^G93A^ mice and patients with ALS (Steyn *et al*., 2020). However, not all patients can switch fuel usage and those who do not switch show signs of faster progression. Fuel usage is also associated with mitochondrial morphology. Mitochondrial fragmentation, which is common in ALS, increases fatty acid oxidation through induction of the enzyme CPT1 (Ngo *et al*). This may, in part, explain the switch to fatty acid dependency seen in some patients with ALS.

We found that the variation in NAT1 expression in ALS was not associated with genetic mutations, suggesting epigenetic regulation. To date, at least three epigenetic pathways have been implicated in NAT1 expression: DNA methylation, histone acetylation, and microRNA expression. Methylation has been reported for both the *NAT1* gene (Kim *et al*, 2008) and *mNat2* gene(Wakefield *et al*, 2010). Hypomethylation results in greater expression. NAT1 transcription is partially repressed by histones and inhibition of histone deacetylases results in increased NAT1 expression (Paterson *et al*, 2011). Finally, several microRNAs including miR-6744-5p (Malagobadan *et al*, 2020), miR-1290 (Choudhury *et al*, 2022; Endo *et al*, 2014) and mir-221 (Guo *et al*, 2020) can decrease NAT1 activity by increasing mRNA degradation. mir-1290 is down-regulated in the motor cortex of ALS patients (Wakabayashi *et al*, 2014) while mir-221 is up-regulated in ALS serum (Taguchi & Wang, 2018). Which of these regulatory pathways is relevant in NAT1 expression in ALS requires additional investigation. If induction of NAT1 at onset is necessary to reduce disease progression, then strategies can be developed to induce NAT1 in the appropriate patient population. For example, the pan-histone deacetylase inhibitor VTR-297 is currently in clinical trials for hematologic malignancies (Przychodzen *et al*, 2022). The current study provides a strong foundation implicating NAT1 in ALS. Further studies in preclinical models of ALS where mNat2 is deleted, or over-expressed, may provide additional insight into the role of NAT1 in disease progression and survival, and whether it is a viable target in metabolic diseases such as ALS.

The heterogeneous nature of ALS has been well documented. In a recent meta-analysis of 76 pre-clinical studies, treatments that target mitochondria in ALS animal models significantly improved survival (Mehta *et al*, 2019). However, translation of these findings to patients has been less impressive (Kiernan *et al*, 2021). Both the heterogeneity and the current etiology of ALS is not well understood. This may contribute to the lack of treatment development. We propose than NAT1 may be a useful biomarker to help distinguish patient sub-populations, especially during clinical trials of new treatments.

## Methods

### Animals

Transgenic mice overexpressing the human SOD1^G93A^ mutation and control animals were purchased from The Jackson Laboratory and bred on a C57 black 6 background. Tissues were collected according to the Animal Ethics Committee (Ethics 2021/AE000152, 2020/AE000094, 2022/AE000707).

### mNat2 qPCR

RNA was extracted from frozen skeletal muscle by grinding in a mortar and pestle on dry ice. RNA was extracted from ∼50 mg of muscle powder using the TRIzol Plus RNA Purification Kit (Invitrogen, cat. no. 12183555) according to manufacturer’s instructions. Briefly, 500 µL TRIzol was added to the muscle powder and homogenised through 19, 22 and 26-gauge needles. An additional 500 µL TRIzol was added, vortexed and incubated for 5 min. Chloroform was added, tubes were shaken vigorously for 15 sec, incubated for 3 min, and centrifuged for 15 min at 4°C and 12,000*g*. The upper phase was transferred to a clean tube and an equal volume of 70% ethanol was added, vortexed and inverted to disperse precipitate. The lysate was transferred to a spin column and centrifuged for 15 sec at 12,000*g*. The column was washed and dried before RNase-free water (30 µL) was added to elute RNA. RNA concentration was quantified by spectrophotometry using the Nanodrop 2000 (Thermo FisherScientific) and samples were stored at -20°C.

First strand complementary DNA (cDNA) was synthesised from 2 µg total RNA in 20 µL reactions using SuperScript IV Reverse Transcriptase (Invitrogen, cat. no. 18090010), as per the manufacturer’s protocol. mNat2 mRNA expression levels were determined by quantitative polymerase chain reaction (qPCR) using the QuantStudio 6 Flex Real-Time PCR System (Applied Biosystems). The cDNA was amplified using primers for mNat2 variant 2 and mGAPDH (see Supplementary Table S5 for details) and the SensiFAST SYBR Lo-ROX Kit (Meridian Bioscience, cat. no. BIO-94005), according to the manufacturer’s protocol. PCR conditions were: 1 cycle for 5 min at 95°C, 35 cycles of: 10 sec at 95°C and 10 sec at 64°C, 1 cycle of 10 sec at 72°C, followed a melt curve. Samples were analysed using the comparative C_T_ method and mNat2 mRNA values were normalised to mGAPDH.

### mNat2 activity

To quantify enzymatic activity, 20 mg of muscle tissue was collected and placed in 2 mL TED buffer (20 mM Tris, 1 mM EDTA, 1 mM DTT, pH 7.4). Tissue was disrupted by sonication with 4 × 5 sec bursts with 30 sec intervals on ice (output = 4; Branson Sonifier 250, Emerson). To pellet cell debris, 1.4 mL lysate was transferred to a tube and centrifuged (Beckman Coulter Microfuge 22R Centrifuge) at 18,000*g* for 10 min at 4°C. The supernatant was transferred to a fresh tube, centrifuged again, and the final lysate was kept on ice. Protein concentrations were assayed by the Bradford method (Bradford, 1976). Enzymatic activity was measured with ∼ 50 µg protein, 440 µM p-aminobenzoic acid (PABA) and 1.1 mM acetyl-coenzyme A for 60 min at 37°C. Reactions were terminated using 25% trichloroacetic acid before neutralising with 1 M Tris-HCl buffer (pH 7.4) and 2 M NaOH. Reactions were centrifuged for 3 min at 15,000*g*. N-acetyl-PABA (NAPABA) was measured using high performance liquid chromatography (Agilent 1100 HPLC Series) as described elsewhere(Butcher *et al*, 2000).

### Subjects

The study was approved by the Royal Brisbane and Women’s Hospital and University of Queensland human research ethics committees. Informed written consent was obtained from all subjects. Experiments conform to the principles set out in the WMA Declaration of Helsinki and the Department of Health and Human Services Belmont Report. For measurements of bioenergetics and NAT1 activity, a total of 33 ALS patients and 24 controls were recruited. Exclusion criteria were a history of any metabolic condition and diabetes mellitus. Body composition was determined using the BodPod system (Cosmed) by whole-body air displacement plethysmography. Resting energy expenditure was measured using a Quark RM respirometer by indirect calorimetry(Steyn *et al*., 2020). Fat free mass was used to predict the resting energy expenditure of the participants as we have done previously(Steyn *et al*., 2018).

### Isolation of PBMC

Human peripheral blood mononuclear cells (PBMC) were isolated by a density gradient centrifugation method using Histopaque-1077 (Sigma-Aldrich, cat. no. 10771) from venous blood anticoagulated with EDTA (Fisher Scientific, cat. no. 02-683-99C). After mixing 2 mL whole blood with an equal volume of PBS, each sample was layered on 4 mL of Histopaque-1077 and centrifuged at 400*g* for 30 min at room temperature. The upper layer was aspirated and discarded, and the opaque interface transferred into a clean tube. The volume was made up to 8 mL with PBS and centrifuged at 300*g* for 10 min. The supernatant was discarded, and the 8 mL PBS wash was repeated. The final PBMC pellet was resuspended in 1 mL PBS for counting and analysis. Cells were frozen in a mixture of 10% DMSO (Sigma-Aldrich, cat. no. D8418), 40% FCS (Fisher Scientific, cat. no. 11950958), and 50% RPMI 1640 medium (Gibco, cat. no. 21870) and stored at -80°C for up to four weeks. Preliminary experiments showed no changes in mitochondrial respiration in cells frozen under these conditions.

### *NAT1* genotyping and sequencing

Genomic DNA (gDNA) was isolated using the ISOLATE II Genomic DNA Kit (Meridian Bioscience, cat. no. BIO-52067), according to the manufacturer’s protocol. DNA concentrations were measured by spectrophotometry using the NanoDrop 2000. The coding region and 3’UTR of the *NAT1* gene was amplified from ∼ 50 ng gDNA using NAT1 specific primers (see Supplementary Table S2 for details) and Platinum SuperFi II Green PCR Master Mix (Invitrogen, cat. no. 12369010), as per the manufacturer’s protocol. PCR conditions were: 1 cycle at 98°C for 3 min, 40 cycles of: 98°C for 10 s, 62°C for 10 sec, 72°C for 2 min, and then 1 cycle at 72°C for 5 min. The PCR product was purified using the PureLink Quick PCR Purification Kit (Invitrogen, cat. no. K310001. Sanger sequencing was conducted with the purified PCR product and BigDye Terminator v3.1 (Applied Biosystems, cat. no. 4337458). PCR conditions were: 1 cycle at 96°C for 2 min and then 30 cycles of: 96°C for 10 s, 50°C for 5 sec, and 60°C for 4 min. Capillary separation was performed by the Australian Genome Research Facility, University of Queensland. The DNA sequences were aligned to a human *NAT1* gene sequence (GenBank accession number NC_000008 with *NAT1*10* allele sequence) to search for any base changes using the National Centre for Biotechnology Information BLAST search engine. Single nucleotide polymorphisms (SNPs) were then used to deduce *NAT1* haplotypes according to the Arylamine N-acetyltransferase Nomenclature database (http://nat.mbg.duth.gr/).

### NAT1 activity

The method detailed above for mouse mNat2 was followed, with minor alterations. PBMC were pelleted by centrifugation at 500*g* for 5 min, and then resuspended in 400 µL TED buffer and lysed by sonication with 3 × 3 sec bursts. The sonication time was reduced to avoid overheating the sample. Protein concentrations were determined, and each reaction used 2-3 µg total protein. NAPABA was measured by HPLC.

### Mitochondrial respiration

Two hours prior to the experiment, the distilled water was aspirated from the XFe96 sensor cartridge utility plate (Agilent Technologies, cat. No. 103022-100) and replaced with 200 µL/well XF calibrant solution. The FluxPak was placed back in the non-CO_2_, 37°C incubator for 45-60 min. In this time, the frozen PBMCs were thawed at 37°C and then centrifuged at 500*g* for 5 min to pellet the cells. The supernatant was discarded, the PBMC were resuspended in 37°C RPMI-1640 complete growth medium (RPMI-1640 medium supplemented with 10% (v/v) FCS, 100 units/mL penicillin and 100 µg/mL streptomycin (Gibco, cat. No. 15140122), and 2 mM L-glutamine (Gibco, cat. No. A2916801)), and incubated at room temperature for 30 min. They were pelleted again resuspended in assay medium at a concentration of 2.75 × 10^5^ cells per 175 µL. Cell suspension was added to each well in quadruplicate in a 96-well Seahorse XF96 Cell Culture Microplate at a density of 2 × 10^4^ cells/well in 80 µL medium. The cell culture microplate was centrifuged at 300*g* for 2 min (Beckman Coulter Allegra X-22R Centrifuge) to bring the cells to the well bottom. The four corner wells contained 80 µL medium only for background correction purposes. The cell culture microplate was then placed in the non-CO_2_, 37°C incubator for 45-60 min. Assay medium consisted of Seahorse XF DMEM medium (pH 7.4; Agilent Technologies, cat. No. 103575-100) supplemented with 1 mM pyruvate, 2 mM glutamine, and 10 mM glucose. During this incubation period, the FluxPak was removed from the incubator and the ports were loaded with compound for final well concentrations of 1 µM oligomycin (port A), 2.5 µM carbonyl cyanide-4-trifluoromethoxyphenylhydrazone (FCCP; port B), and 1 µM rotenone/antimycin A (port C). It was then loaded into the Seahorse Xfe96 Analyzer (Agilent Technologies) and calibrated. Once the calibration was complete, the utility plate was replaced with the cell microplate and the experiment was started. The Wave software (Agilent Technologies) pre-set protocol for the Mito Stress Test were used for timed, sequential injections of oligomycin, FCCP, and rotenone/antimycin A at 16, 36, and 56 mins, respectively. The mean of the three measurements after each injection, per well, was used for the calculation of basal respiration, ATP production, proton leak, maximal respiration, non-mitochondrial respiration, reserve respiratory capacity, basal glycolysis, glycolytic reserve, and maximal glycolysis.

### Statistical analysis

GraphPad Prism 9 was used for all statistical analysis. For one-way analysis of variance, the significance level was adjusted for multiple comparisons using Bonferroni’s test. Significant differences between two groups were determined using Mann-Whitney test assuming significant if p < 0.05. The association between two variables was determined using Spearman’s rank order correlation and denoted by rho (ρ). Data are presented as mean ± standard error of the mean (s.e.m.). For patient studies, a power calculation was performed to determine the minimum number of subjects required to achieve an 80% power (beta = 0.2) with a type I error of 0.05. Using historical data for PBMC respiration where standard deviation is approximately 50% of the mean, a minimum of 19 independent samples in each group was required to show a 45% difference between measurements in controls and patients.

## Acknowledgements

The authors thank the contributions from the control volunteers and ALS patients who generously provided blood samples for this study. The authors also acknowledge funding from the National Health and Medical Research Council of Australia (grant 1083036). CC is the recipient of a Commonwealth Government Postgraduate Scholarship.

---

**Supplementary Table S1.** – Normality tests.

| Parameter | D'Agostino & Pearson test<br>(p value) |  | Shapiro-Wilk test<br>(p value) |  |
| --- | --- | --- | --- | --- |
|  | Controls | Cases | Controls | Cases |
| Basal respiration | 0.63 | 0.0002 | 0.69 | 0.0013 |
| ATP production | 0.52 | 0.001 | 0.34 | 0.002 |
| RRC | 0.033 | 0.047 | 0.029 | 0.0018 |
| Max. respiration | 0.067 | 0.023 | 0.054 | 0.0022 |
| Proton Leak | 0.34 | <0.001 | 0.075 | <0.001 |
| Non-mito respiration | 0.09 | 0.025 | 0.148 | 0.025 |
| Basal Glycolysis | 0.68 | 0.18 | 0.65 | 0.19 |
| Glycolytic reserve | 0.38 | 0.07 | 0.17 | 0.055 |
| Maximal glycolysis | 0.21 | 0.17 | 0.45 | 0.12 |
| Metabolic potential | 0.033 | 0.048 | 0.028 | 0.002 |
| NAT1 | 0.07 | 0.43 | 0.15 | 0.92 |

**Supplementary Table S2.**
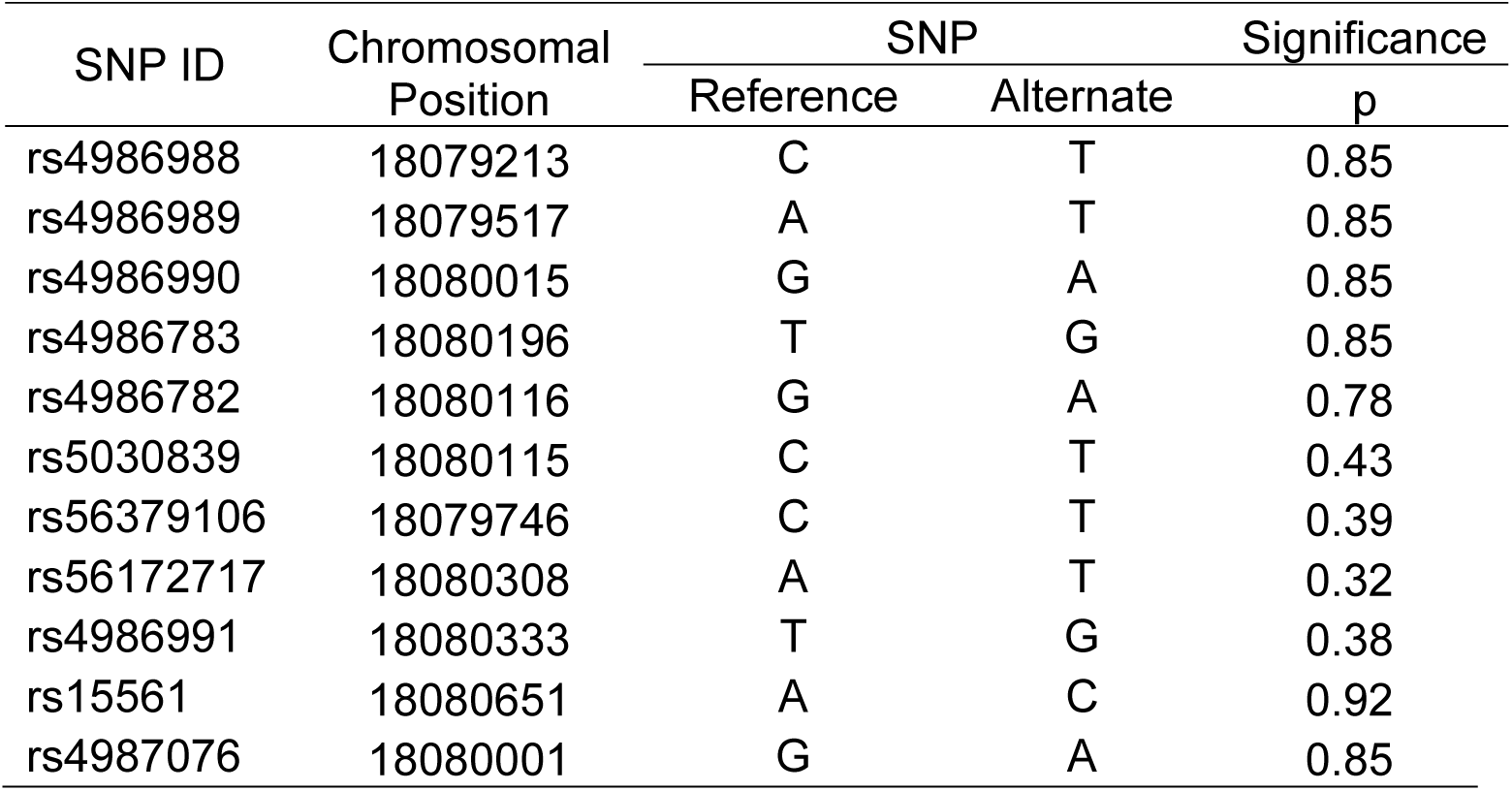
NAT1 single nucleotide polymorphisms in ALS.

| SNP ID | Chromosomal<br>Position | SNP |  | Significance |
| --- | --- | --- | --- | --- |
|  |  | Reference | Alternate | p |
| rs4986988 | 18079213 | C | T | 0.85 |
| rs4986989 | 18079517 | A | T | 0.85 |
| rs4986990 | 18080015 | G | A | 0.85 |
| rs4986783 | 18080196 | T | G | 0.85 |
| rs4986782 | 18080116 | G | A | 0.78 |
| rs5030839 | 18080115 | C | T | 0.43 |
| rs56379106 | 18079746 | C | T | 0.39 |
| rs56172717 | 18080308 | A | T | 0.32 |
| rs4986991 | 18080333 | T | G | 0.38 |
| rs15561 | 18080651 | A | C | 0.92 |
| rs4987076 | 18080001 | G | A | 0.85 |

**Supplementary Table S3.**
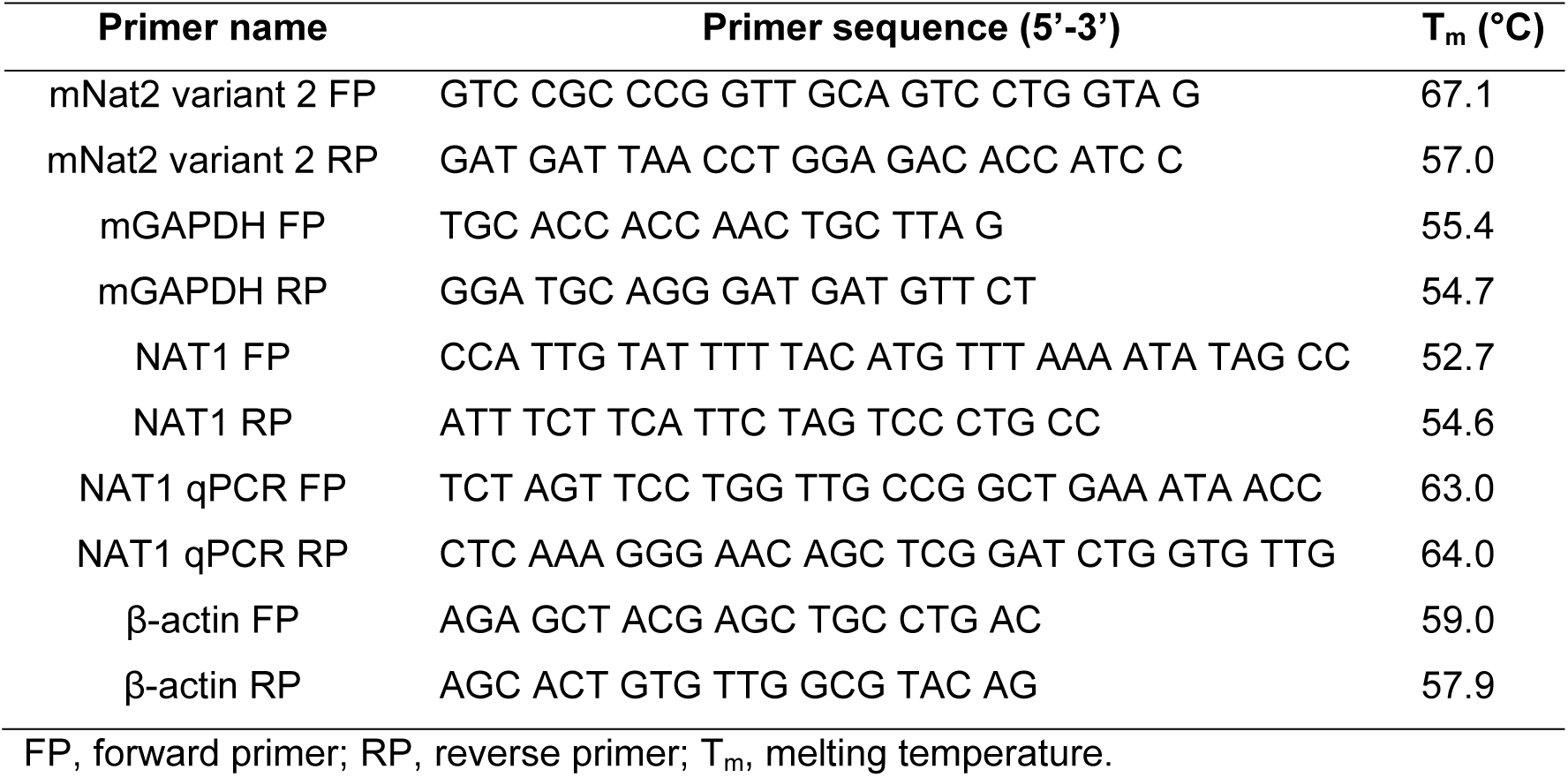
– PCR primers.

